# Machine Learning Approaches Identify Genes Containing Spatial Information from Single-Cell Transcriptomics Data

**DOI:** 10.1101/818393

**Authors:** Phillipe Loher, Nestoras Karathanasis

## Abstract

**Motivation:** We participated in the DREAM Single Cell Transcriptomics Challenge. The challenge’s focus was two-fold; a) to identify the top 60, 40 and 20 genes that contain the most spatial information, and b) to reconstruct the 3-D arrangement of the D. melanogaster embryo using information from those genes.

**Results:** We developed two independent approaches, leveraging machine learning models from Lasso and Deep Neural Networks, that we successfully apply to high-dimensional single-cell sequencing data. Our methods allowed us to achieve top performance when compared to the ground truth. Among ~40 participating teams, the resulting solutions placed 10th, 6th, and 4th in the three DREAM sub-challenges #1, #2 and #3, respectively. Notably, for the Lasso approach we introduced a feature selection technique, Lasso-TopX, that allows a user to define a specific number of features they are interested in and the Neural Network approach utilizes weak supervision for linear regression to accommodate for uncertain or probabilistic training labels. Furthermore, we identified novel *D. melanogaster* genes that carry important positional information and were not previously suspected. Lastly, we show how the indirect use of the full datasets’ information can lead to data leakage and generate bias in overestimating the model’s performance.

**Availability:** https://github.com/TJU-CMC-Org/SingleCell-DREAM/.

## Introduction

Single-cell RNA sequencing (scRNA-seq) has been rapidly gaining popularity and allows biologists to gain knowledge about the abundance of genes for thousands of cells, individually, from a given tissue. Such an approach does not suffer from the drawback of standard approaches where the aggregation of a large starting population of cells obscure the ability to detect cell-to-cell variation. Unfortunately, scRNA-seq approaches do not typically maintain the spatial arrangement of the cells (Karaiskos *et al.*, 2017).

For more than a decade, the DREAM (Stolovitzky *et al.*, 2007) initiative has driven crowd-sourced open science scientific contests in different areas of biology and medicine. Recently, the DREAM Single Cell Transcriptomics Challenge (Tanevski *et al.*, 2019), in which we participated, focused on tackling the reconstruction of the 3-D arrangement of cells using predefined number of genes. Specifically, the goal of this DREAM challenge was to use *D. melanogaster* embryo as a model system and seek to determine whether one can reconstruct the spatial arrangement of cells from a stage 6 embryo by only using a limited number of genes. The challenge piggy backed off previously published scRNA-seq datasets and a computational mapping strategy called DistMap, that leveraged in-situ hybridization data from 84 genes of the Berkeley Drosophila Transcription Network Project (BDTNP), which was shown to uniquely classify almost every position of the *D. melanogaster* embryo (Karaiskos *et al.*, 2017). Out of these 84 genes (herein referred to as “inSitu genes”) and without using hybridization data, the participants were asked to identify the *most informative* 60, 40, and 20 genes for subchallenges #1, #2, and #3 respectively. In addition to gene selection, each subchallenge also required participants to submit 10 locations predictions (X, Y, Z coordinates) for each of the cells using only the selected genes (Tanevski *et al.*, 2019).

In order to identify the most informative genes, we describe two independent feature selection strategies. The first, we name Lasso-TopX, leverages linear models using the Least Absolute Shrinkage and Selection Operator (Lasso) (Friedman *et al.*, 2010; Tibshirani, 1996). Lasso has a few important characteristics that made it desirable to use. Specifically, the models are easy to interpret because each feature gets assigned a coefficient and the coefficients are combined linearly. It is also useful for dimensionality reduction because the resulting coefficients can be exactly zero, essentially eliminating features (James et al., 2013; Friedman et al., 2010). Our second feature selection strategy leverages Deep Neural Networks (NN). NNs are making major advances in problem solving by allowing computers to better discover structure in high-dimensional data (LeCun *et al.*, 2015). By linking multiple non-linear layers together, we sought to use Deep Learning in order to discover subsets of genes that would not have otherwise been possible with more traditional linear approaches.

In what follows, we describe our methods and the novel elements that allowed us to meet the objectives of the DREAM challenge. Notably, Lasso-TopX allows a user to specify the exact number of key features they are interested in. And to take advantage of Distmap’s probabilistic mapping where a cell’s location is not always unique, we also describe how NNs can be trained using weak supervision (Zhou, 2018) for use in linear regression. Importantly, while not an objective of the DREAM challenge, we extend our techniques to other genes by looking for *non*-inSitu genes that also carry spatial information.

## Methods

In summary, we used two methodologies to identify the most informative features (*D. melanogaster* genes); an approach based on Deep Neural Networks (NN) models, and an approach based on Lasso models, which we call Lasso-TopX. Both are supervised approaches that use training data. We then utilized inference techniques on the trained models to get a list of the most important 60 / 40 / 20 inSitu genes. In order to help baseline our results prior to the end of the competition, we also leveraged a process (herein named Random) that picked genes randomly. For the selected genes using NN, Lasso-TopX, and Random, we passed only those genes into DistMap (Karaiskos *et al.*, 2017) to get the spatial predictions.

### Data made available by competition organizers

Below is a summary of the data provided to us by the DREAM challenge:

- *Reference database:* The reference database comes from the expression patterns (Fowlkes *et al.*, 2008) of the in-situ hybridizations of 84 genes from the Berkeley Drosophila Transcription Network Project project. The *in-situ* expression of 84 genes is quantified across the 3039 *D. melanogaster* embryonic locations.
- *Spatial coordinates:* X, Y, and Z coordinates were supplied for the 3039 locations of the *D. melanogaster* embryo.
- *Single cell RNA sequencing:* Three expression tables were provided; the raw, normalized, and binarized expression of 8924 genes across 1297 cells (Karaiskos *et al.*, 2017).
- *DistMap source code* was provided and it was used to identify the cell locations in the initial publication (Karaiskos *et al.*, 2017).

### Cell Locations

For each cell (n=1297) available in the RNA sequencing data, we generated training labels representing their 3-D positions by running DistMap (Karaiskos *et al.*, 2017) with the following inputs:

- single cell RNA sequencing expression data, both raw and normalized
- the Reference database
- the spatial coordinates

Briefly, DistMap calculates several parameters, a quantile value and one threshold per inSitu gene, to predict the cell locations. It employs these values to binarize the expression of the genes’ and calculate the Matthews Correlation Coefficients (MCC) for every cell-bin (embryo location) combination. By doing this, DistMap maps a cell to multiple likely positions. Lasso-TopX and NN approaches (described below) use these MCC values to determine the training labels.

### Feature selection approaches

#### Random

As a baseline approach, among the 84 inSitu genes from which we were allowed to pick, we randomly selected 60, 40 and 20 of them for the respective subchallenges. This random selection allowed us to benchmark (see Results) how Lasso-TopX and NN feature selection approaches compared against a random process. We performed this selection step 10 times, one for each outer cross validation fold, see “Post Challenge Outer Cross Validation” section, below. The importance of this comparison is to evaluate if the cost of building a method, both timewise and computationally, has any advantage over a simple approach, that does not leverage machine learning (Karathanasis *et al.*, 2014).

#### Lasso-TopX

We introduce a method, Lasso-TopX, that is implemented in the R programming language and leverages the glmnet package (Friedman *et al.*, 2010) to build generalized linear models with Lasso (Tibshirani, 1996). This method allows for the identification of the most informative N features, where N is 60, 40, and 20 for sub challenges 1, 2, and 3 respectively.

##### PRE-PROCESSING

We used the following data to identify the most important genes:

- *Single cell RNA sequencing*: we subset the provided normalized single cell RNAseq dataset to include only the 84 *in-situ* genes.
- *Top cell locations*: for training labels, we identified the locations of the cells using distMap with the code provided from the challenge’s organizers, see section ‘Cell Locations’ above. For each cell, we use the bin (embryo locations) corresponding to the maximum MCC. In our feature selection process, we employed only the cells that are mapped uniquely to one location (1010 out of 1297 cells), Supplementary Figure S1.

##### TRAINING FLOW AND FEATURE SELECTION

We performed the following steps to identify the important features employing Lasso-TopX:

1. In order to identify the most important 60 / 40 / 20 features we performed a repeated five-fold cross validation (CV) process. The CV was repeated 20 times for 300 different values of lambda. Lambda is Lasso’s hyperparameter which the user needs to optimize. Intuitively, fewer features will be selected as lambda increases. The range of the lambda values was defined manually, using 70% of the data and only one time, in order for models with 60 / 40 / 20 features to be produced. In relation to the competition, we retrieved lambda ranges from glmnet packages, 100 values, and tripled the density to include 300 values. In total, we fitted 5*20*300 = 30,000 models. Importantly, in order to avoid overfitting, during each CV fold only the training data corresponding to this fold are standardized and the resulting model is applied to the test data (Friedman *et al.*, 2010).
2. For each model, we extracted the following information
  a. *The error from the model*: the Euclidean distance of the predicted XYZ location to the top location.
  b. The number of features which were used and their corresponding coefficients.
3. We selected the *best lambda* value by calculating the mean error per lambda across the repeated five-fold CV (Figure 1). The *best lambda* was producing the minimum mean error and models with the desired number of features. We retained only the models corresponding to this lambda value. One lambda value and one hundred models (5-fold CV * 20 times) were selected per sub-challenge.
4. For each one of the selected models we extracted their features and calculated two metrics.
  a. Stability: the number of times a feature was selected as important across the repeated cross validation procedure (Figures 1b left, for sub-challenge 3), and
  b. Mean Coefficient: the mean value of the coefficients that a feature was assigned across all coordinates (Figure 1b right), for sub-challenge 3.
5. Finally, we utilized the RankSum statistic to combine these two metrics and calculate the overall importance of the selected features. In cases where we had more than the desired number of features in our final feature list we kept the features with the higher RankSum statistic (Figure 1). Having more than the desired number of features is possible as models with the same lambda may select different features during the repetitive cross validation process.

**Figure 1.**
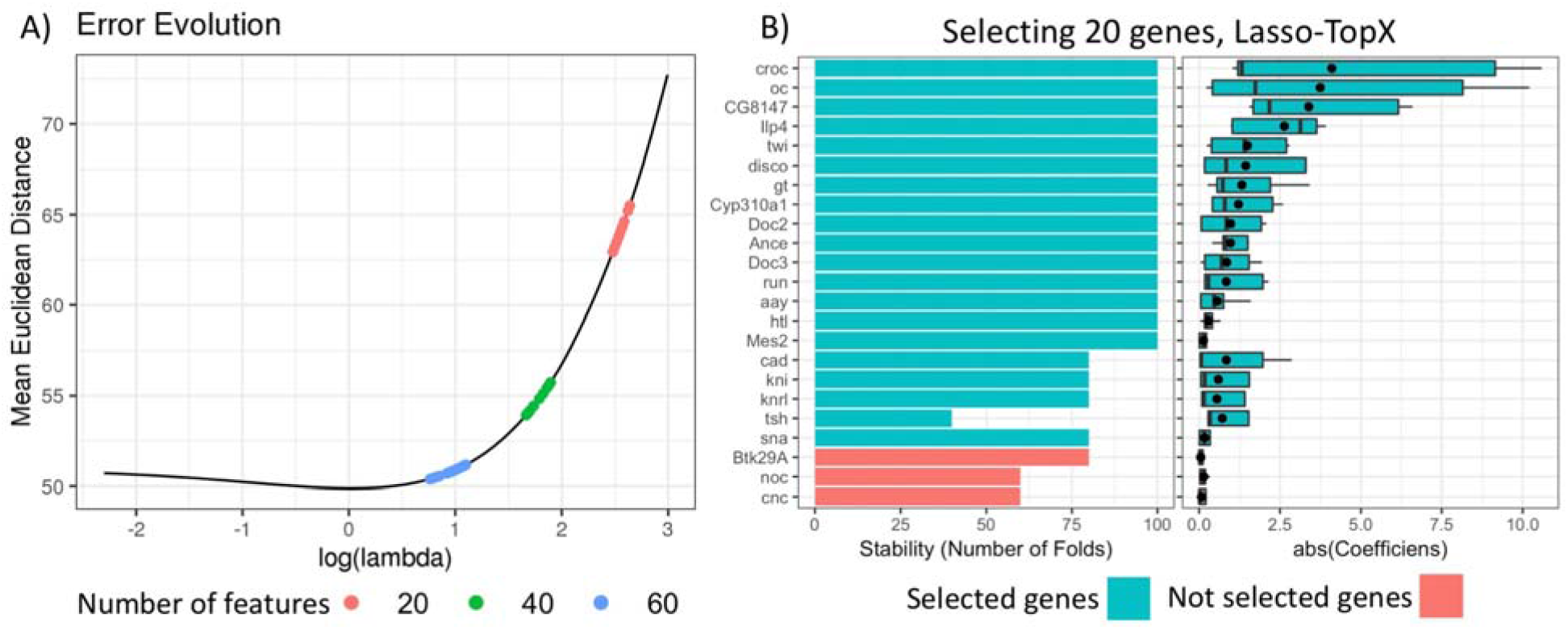
Selecting 20 genes – Lasso-TopX. **A) Lambda Selection:** The mean Euclidean distance error across the repeated fivefold CV in relation to the natural logarithm of lambda is presented for one out of 10 outer Cross Validation runs. Several models with 20 features (red dots) are produced with their lambda values to range from 14.00083, log(14.00083) = 2.64, to 11.93777, log(11.93777) = 2.48. The lambda that produced the models with the minimum Mean Euclidean Distance error was selected. The same process was followed for both 40 (green dots) and 60 (light blue dots) features. **B) Left):** Stability, the number of times a feature was selected across the repeated cross validation procedure. 20 genes are shown in one out of the 10 outer Cross Validation runs. Each gene was selected at least one time by the best performing models during the repeated cross validation (lambda = 11.93777). 20 genes with the higher RankSum statistic were selected (Blue), the last three genes (red), had the lowest RankSum statistic and left out from the final list. **B right)** The distribution of the absolute value of the coefficients of the selected genes. The mean value of the coefficients is shown with a black point character.

### Neural network-based approach using weak supervision

In this approach, we perform weakly supervised learning (Zhou, 2018) using Neural Networks. After training the models, we calculate variable importance scores to rank each gene. We describe several techniques that we used to help eliminate overfitting and make sure our model generalizes well. Because the training labels were not given to us directly and because we could not assume the max MCC from DistMap was always correct, we devised a technique that is able to use multiple training labels for the same set of input neuron values.

#### PRE-PROCESSING

All genes (n=8,924) from the normalized RNAseq dataset, ‘dge_normalized.txt’ were used as predictor variables. For generating the training labels for the 1,297 cell locations, we used the MCC based procedure also used by Lasso-TopX, but with one modification. Instead of using only the location (X, Y, Z) from the max MCC score, we used all locations that had an MCC score greater or equal than 95% of the max MCC score.

#### NN ARCHITECTURE AND LOSS FUNCTION

Model training and inference were all performed using Python 3.5, the PyTorch (Paszke, Adam, Sam Gross, Soumith Chintala, Gregory Chanan, Edward Yang, Zachary DeVito, Zeming Lin, Alban Desmaison, Luca Antiga, 2017) machine learning library, and the numpy and pandas packages. We used a fully-connected NN architecture described below.

- *Input neurons*: one input neuron per RNAseq gene
- *Hidden layers*: two hidden layers were used, each with 100 neurons
- *Output neurons*: three output neurons were used (X position, Y position, and Z position)
- *Loss function*: Euclidean Loss using all three output neurons
- *Activation function*: Rectified Linear Units (Nair and Geoffrey E., 2017; Hahnioser *et al.*, 2000)
- *Optimizer*: Adadelta (an adaptive learning rate method) was used for gradient descent (Zeiler, 2012)
- *Other*: To help avoid overfitting and allow the model to generalize better, we used Hinton Dropout (Srivastava *et al.*, 2014), set at 10% for both the hidden and input layers.

#### TRAINING FLOW

Using the pre-processed data, we performed 5-fold cross validation 40 times for a total of 200 models. Before training, the genes were first standardized to have a mean of 0 and unit variance. For each model and to prevent bias (Ambroise and McLachlan, 2002), the parameters used for this adjustment were determined from the training splits only, and then applied to the validation split.

Similarly, using the training split only to avoid selection bias (Ambroise and McLachlan, 2002; Smialowski *et al.*, 2009), we removed correlated genes by first identifying genes that (1) were not an inSitu-gene and (2) had a Pearson correlation with at least one inSitu-gene of >= 0.6 or <= −0.6. We then removed these RNAseq genes from all splits. This allowed us to remove non-inSitu genes that might prevent variable importance scores of inSitu genes from showing up which we thought might be helpful because the subchallenges only asked us to report inSitu genes.

At every 80/20 split, we made sure that a cell’s gene expression values were never found in both the training and validation splits. During training, minibatch sizes of 100 were used. To help prevent the model from overfitting on the training data, we also performed early-stopping by stopping the training process once the Euclidian loss on the validation fold did not improve after 50 epochs and keeping the model with the lowest loss. Because looking at the validation fold’s loss function could potentially overestimate a model’s performance, we evaluated our subchallenge scores on an external hold-out set from a separate outer cross-validation fold (see Results).

#### FEATURE SELECTION

Variable importance (VIP) scores were calculated and ranked for each of the 200 NN models used in the training process. We implemented (and include in our source code) Gedeon’s method (Gedeon, 1997) to come up with Variable Importance Scores for each model. For each model, we only kept the genes with the highest 60/40/20 VIP scores depending on the subchallenge. We then sorted the lists by consensus vote to get one list per subchallenge. For the subchallenges, only inSitu-genes were selected.

### Location predictions

The following steps were used by Lasso-TopX, NN and Random approaches in order to predict 10 locations per cell:

1. We subset the *in-situ* expression database, keeping only the 60, 40, 20 genes as they were identified by the feature selection steps above.
2. We calculated MCC applying DistMap with the following modifications:

2.1. We employed only the cells belonging in the training sets and the selected genes to calculate DistMap’s parameters, which are used to binarize the genes’ expression data (Karaiskos *et al.*, 2017).
2.2. We used the same parameters to binarize the expression data of the cells belonging in the test set.
2.3. Finally, similar to “Cell Locations” section above, we calculated the MCC for every cell-bin combination and we selected the 10 bins that correspond to the top 10 highest MCC scores.

### Post Challenge Outer Cross Validation

In the post-challenge phase, the organizers split the data in 10 folds, on which our approaches were re-run for stability and overfitting evaluation (Tanevski *et al.*, 2019). Separately for each of the 10 iterations, only the respective 9 training folds were used for feature selection and to train Distmap. The cells’ locations were predicted for the remaining validation fold. We refer to this post challenge cross validation as the ‘outer cross validation’ because any cross-validations described in our feature selection methods occurred using data only *within* the training-folds of this outer cross-validation.

### Blind evaluation metric

Prior to the competition ending, in which contestants did not have access or insight into the challenge organizers’ scoring functions, we evaluated our location predictions by calculating for each cell the mean Euclidean distance of the top 10 predicted locations from the cell location with the maximum MCC (*MeanEuclDistPerCell*). For the cells that did not map uniquely, we used the first bin among the ties as returned by R.

Then, we calculated the mean of the MeanEuclDistPerCell per outer cross-validation fold across all cells which we refer to as *MeanEuclDistPerFold*. Finally, the mean of the *MeanEuclDistPerFold* across all 10 outer cross-validation folds was calculated and referred to as *MeanEuclDistAllFold*.

## Results

### Challenge submission (Lasso-TopX or NN)

We ran both the Lasso-TopX and NN approaches for all three subchallenges. Because the challenges final scoring algorithms were not available to any participants until after the competition concluded, we compared our two feature selection approaches using a blind evaluation metric (see Methods) we devised and thought might be a proxy to a good leadership score. Because the teams were only allowed one final submission to each subchallenge, we used this blind metric to determine if the results from either NN or Lasso-TopX would be submitted to each subchallenge. This blind metric was calculated individually for both feature selection methods and for each subchallenge. Our evaluation metric suggested that Lasso-TopX may perform slightly better than NN for some subchallenges (data not shown). Based on this, our final submission used results based on NN for subchallenge 2 and Lasso-TopX for the other two. Our submitted results ranked 10th, 6th, and 4th in the three sub-challenges, respectively, among ~40 participating teams (Tanevski *et al.*, 2019).

### Evaluation Post-Challenge

After the challenge ended, the organizers devised a post-challenge cross-validation scheme (see Methods and (Tanevski *et al.*, 2019) for more detail) to evaluate the robustness of the methods. It was only after this resubmission phase did the organizers make the true scoring functions (“s1”, “s2” and “s3” scores) publicly available. Supplementary Figure S2 and Figure 2 show the results of our blind, s1, s2, and s3 metrics across the outer 10-fold cross-validation. As expected, both Lasso-TopX and NN behaved better than Random. The results of the three scoring schemes (Figure 2) agreed with our previous findings using our blind metric, and in agreement with the challenge paper (Tanevski *et al.*, 2019) show that our scores have little variability and that our methods generalize well. Because of the higher s2 score in subchallenge #1 for NN (Figure 2), we note the possibility that our NN approach could have ranked more favorably in subchallenge #1 when compared to the submitted Lasso-TopX predictions.

**Figure 2.**
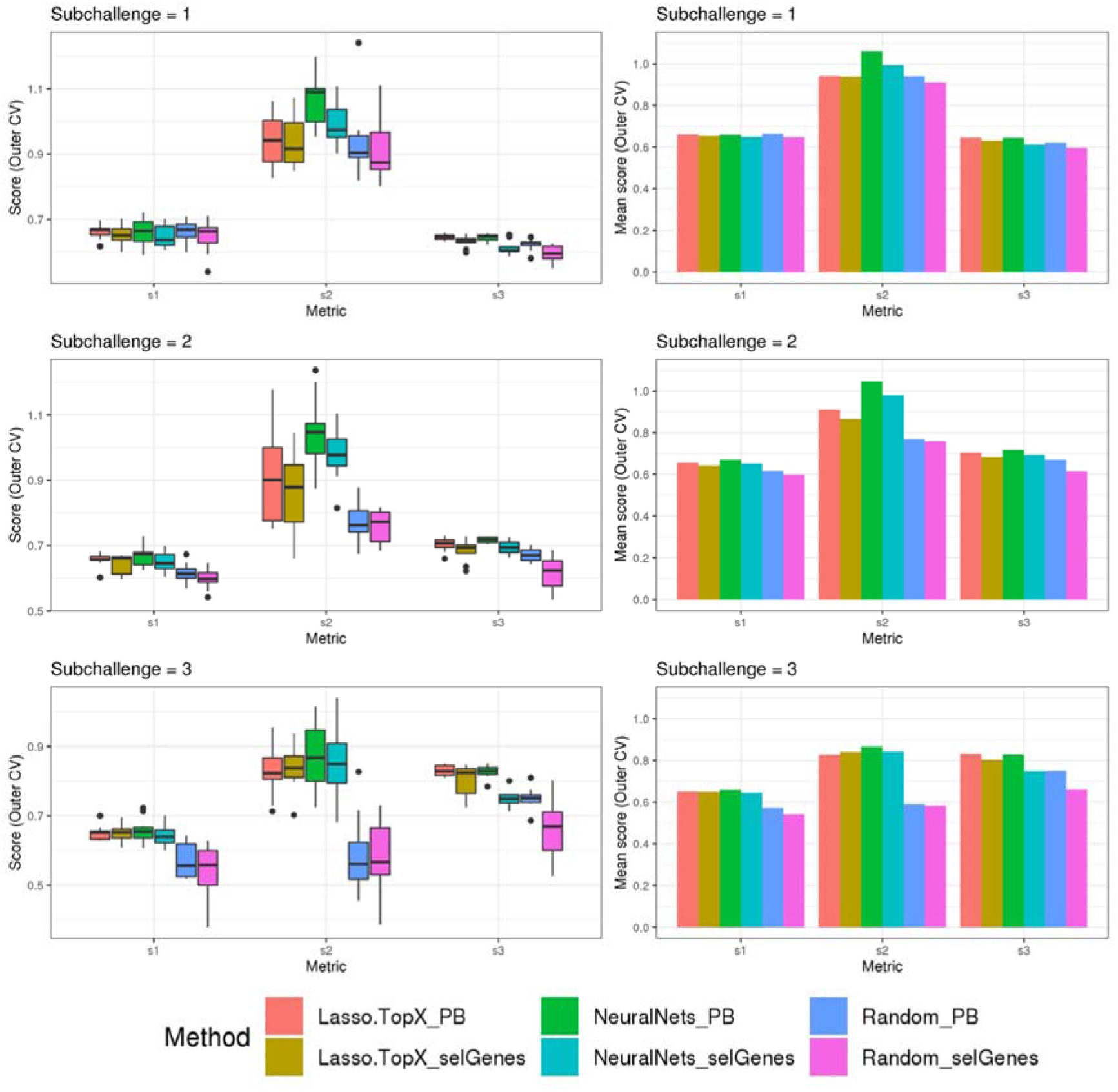
Comparison of different methods, organizers scoring functions. Lasso.TopX (Lasso.TopX_selGenes), performed better for sub-challenges 1 and 3, and NeuralNets (NeuralNets_selGenes) performed better for sub-challenge 2. For sub-challenge 1, Lasso.TopX performed better than NeuralNets for s1 and s3. For sub-challenge 2, NeuralNets performed better for all scores and for sub-challenge 3 Lasso.TopX performed better for s1 and s3 scores. In all sub-challenges, both methods performed better than Random (Random_selGenes). The binarized expression data that were produced using all expression data, _PB extension, showed an extreme bias in overestimation of performance, across all metrics, methods and sub-challenges. Lasso.TopX_PB performed always better than Lasso.TopX_selGenes, NeuralNets_PB performed always better than NeuralNets_selGenes and Random_PB performed always better than Random_selGenes.

### Considerations for Weak Supervision

When generating training labels during the pre-processing step of NNs (see Methods), there were several reasons why we allowed multiple training labels for the same cell. First, it allowed all locations of the 287 cells (Supplementary Figure S1) that did not uniquely map to be used during training. Also, we could not be certain that the max MCC was always the right value to use and wanted to better leverage the probabilistic mapping strategy enabled by DistMap.

Figure 3a shows that the vast majority of the time, there exists more than one spatial location for a cell when using the 95% cutoff. The most common number of selected training labels per cell location is 5, with a mean of 7. When allowing multiple training labels per cell, our dataset became much larger: 11,491 observations instead of only 1,297 observations when only the value with the max MCC was used. One pleasant consequence of having more training data is that it makes it harder to overfit a neural network which is especially problematic in high-dimensionality settings (Verleysen *et al.*, 2003; Bellman, 1961).

**Figure 3.**
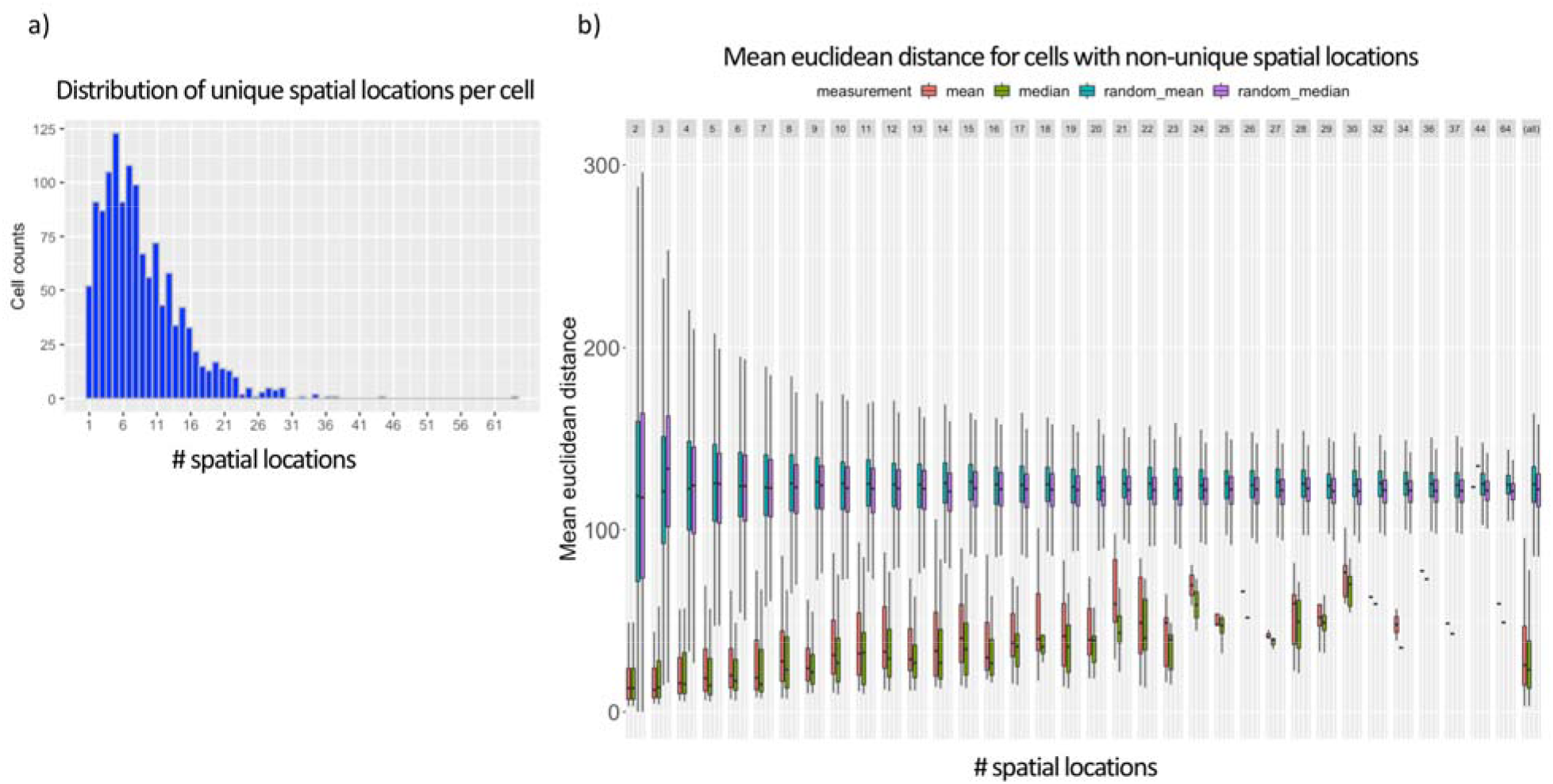
Ambiguous training labels leveraged in NN approach. (a) Shows that most D. melanogaster cells contain more than one spatial location (mean of 7) when using DistMap’s predictions and thresholding at 95% of the max MCC. (b) Shows the mean and median Euclidean distances between the cell’s with non-unique training labels that meet the same 95% threshold. Cell’s that are selected at this threshold have mean and median Euclidean distances much lower than random and highlight the probabilistic nature of the training data.

In Figure 3b we also show that the surviving training labels per cell generally represent similar spatial coordinates when compared with randomly shuffled locations in the training data. This suggested that allowing multiple training labels per cell during training could guide the model to generalize to less-specific spatial regions without being pegged to any one location that could have been incorrectly classified.

Importantly, as mentioned in Methods, when generating training and validation sets we made sure that a cell’s gene expression values were never found in both sets. We accomplished this by splitting on cell names versus the row indices. This is especially important because the pre-processing steps allowed the same cell to be found multiple times but with different training labels (Figure 3a). We found that if we did not split this way, we would have indirect data leakage (Luo *et al.*, 2016) and significantly overestimate the performance of our models because the validation split could contain identical predictor variables (gene expression levels) as the training splits but with training labels that had similar (though not identical) spatial locations (Figure 3b).

### Measuring and avoiding data leakage during location prediction

We also sought to determine what our scores would have looked like if data leakage occurred during the location prediction stage. In machine learning and statistics, *data leakage* can lead to inflated performance estimates when data from the validation or test set are used during training (Luo *et al.*, 2016). Overfitting because of data leakage would have been easy to do by mistake because the provided binarized expression data, generated by DistMap, were produced using all expression data and consequently should never be used at any step of training or testing. For example, one might think that instead of modifying DistMap to perform the two-step approach described in Methods, a contestant could have used the provided binarized data to directly calculate the MCC scores and the 10 cell positions. However, as is evident from Supplementary Figure S2 and Figure 2 this will lead to overestimation of performance irrespective of the scoring functions (blind, s1, s2, s3) or the methods (NNs, Lasso-TopX, Random) used. In both figures we present bars and boxplots which correspond to the overfitted location predictions using the unmodified and provided binarized data (extension “PB”) and compare it to the approach we used (extension “selGenes”).

### InSitu Genes with Spatial Information

We observed that the genes selected across the outer 10 cross validation folds were stable, (Supplementary table S1). More specifically, in the Lasso-TopX case, 74, 54 and 27 genes were selected in total for sub-challenge 1, 2 and 3, respectively, with 44 (60%), 25 (46%) and 15 (56%) of them to be selected across all folds, Supplementary Figure S3a. Similarly, in the NNs case, 64, 47 and 23 genes were selected, with 58 (91%), 33 (70%) and 13 (57%) of them to be selected across all folds, for sub-challenge 1, 2 and 3, respectively (Supplementary Figure S3b). As expected, Random did not show the same trend (Supplementary Figure S3c) with zero genes selected across all folds. We observed agreement in the in-situ genes selection between our two distinct feature selection strategies despite differences in pre-processing, features used during training, and inference models. Specifically, we observed a mean of 80%, 76% and 64% agreement for sub-challenge 1, 2 and 3 respectively, across the outer cross validation folds (Supplementary Figure S4).

### Non-inSitu Genes with Spatial Information

While not a focus on this competition, we additionally ran both our methodologies using RNASeq data information from *both* the non-inSitu and inSitu genes and we were able to discover many informative non-inSitu genes that also contain positional information. The list of 20/40/60 genes using the NN methodology would have been composed of 56%, 50%, 52%, respectively, of non-inSitu genes (Supplementary table S1). In the Lasso-TopX case, 67% 66% and 70% of the selected genes were non-inSitus when selecting the most informative 20 / 40 / 60 genes (Supplementary table S1). Similar to the inSitu genes analysis, we calculated the stability of the selected genes across the outer cross validation folds. In the Lasso case, 36.6%, 28.7% and 23.7% of genes were selected across all folds of the outer CV, when selecting for 20, 40 and 60 genes, respectively (Supplementary Figure S5a). In the NNs case, 46%, 59% and 67% of genes were selected across all folds when selecting 20, 40 and 60 genes, respectively, Supplementary Figure S5b. Furthermore, we observed that on average 52%, 57% and 51% of genes identified by NN and Lasso-TopX were in common across the 10-fold outer cross validation folds, when selecting 60, 40 and 20 genes, respectively, (Supplementary Figure S4b). Interestingly, we observed (Figure 4) that several non-inSitu genes were selected consistently across all 60 feature selection runs (60 = 2 methods * 3 subchallenges * 10 outer CV folds). Specifically, 142 genes were identified across all runs consisting of 38 inSitus and 104 non-inSitus. As expected, due to the fact that inSitu genes contain spatial information, they were selected on average more often, 25 times out of 60, than non-inSitus, 14 out of 60. However, focusing on the most stable genes, genes that were selected in at least 30 out of the 60 runs, 15 out of 29 are non-inSitus, (Figure 4).

**Figure 4.**
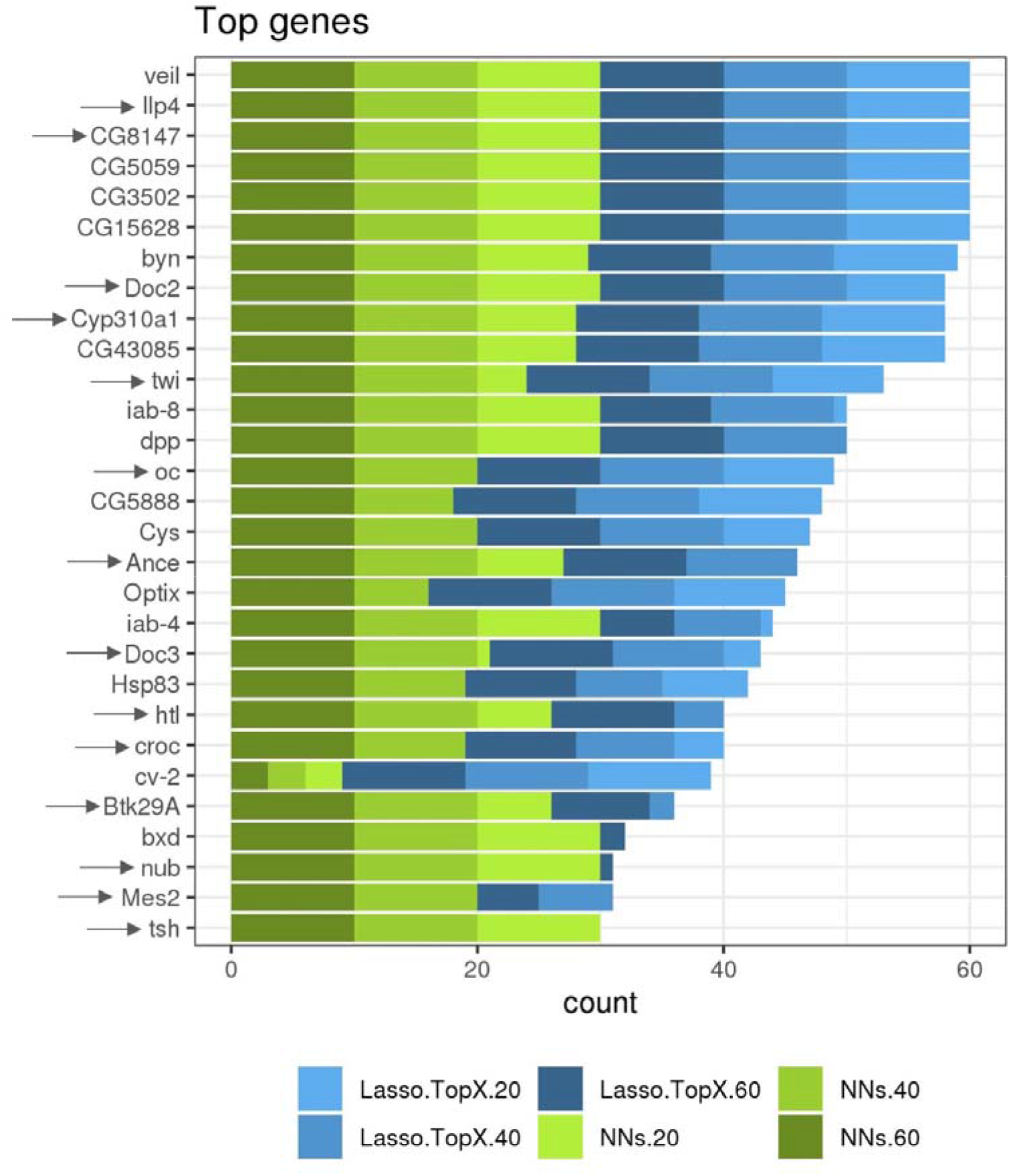
More frequent identified genes. Several inSitu and non-inSitu genes were selected across the sub-challenges, methods and outer cross validation folds. 29 genes, 14 inSitu(arrow) and 15 non-inSitu were selected in at least 30 out of a total of 60 feature selection runs, xaxis. Lasso-TopX and NNs are shown in shades of blue and green, respectively. Dark, regular and light shades correspond to selecting 60, 40 and 20 genes, respectively.

## Discussion

In this paper we present a modified Lasso workflow, named Lasso-TopX, that is able to extract the most important user defined number of features and a NN approach that uses weak supervision that is able to use multiple training labels per cell for linear regression. We developed these methods as part of our participation in the DREAM single cell transcriptomics challenge (Tanevski *et al.*, 2019). The challenge had two main objectives, a) to identify the top 60, 40 and 20 genes that contain the most spatial information, and b) to reconstruct the 3-D arrangement of the *D. melanogaster* embryo using information from those genes. The 20 and 60 genes were identified using Lasso-TopX and the 40 genes using NN. In all cases, the 3-D locations were predicted using Matthew Correlation Coefficients based methodology. Our team, named DeepCMC, ranked 10th, 6th, and 4th in the three sub-challenges, respectively, among ~40 participating teams from all around the world (Tanevski *et al.*, 2019).

A typical Lasso workflow consists of first identifying the best lambda, using cross validation and then employing that lambda to train on the full dataset to identify the most informative features (James et al., 2013). This process does not allow the user to specify a discrete number of features they are interested in because the selected lambda is not tied to a user-defined number of features (Friedman *et al.*, 2010). Also, running the typical workflow multiples times could lead to slightly different optimal lambda values as the data splits during the cross validation could differ and thus this could lead to slightly different features and different number of features. Another approach that could be employed to meet the subchallenges requirements, is to have performed the typical workflow and then select the top 20 / 40 / 60 genes from the resulting list of genes. This approach is not optimal; for example, if someone could select the best 20 features, these 20 features would not necessarily be a subset of the best 40 features (James et al., 2013).

Considering the above, we developed Lasso-TopX, that leverages Lasso and is able to identify the most important user-defined number of features, employing repeated cross validation to make the results less dependent on any particular choice of data split. Lasso-TopX can be applied to classification or regression problems where finding the most important, stable, and user-defined number of features is important.

We also note that Lasso-TopX will provide the most value if the user-defined number of features is less than what the traditional Lasso workflow would have chosen. Taking as an example, figure 1A, we see that Lasso’s error is decreasing in the beginning as we move to higher values of lambda (from left to right), then there is a local minimum (close to log(lambda) = 0), and then the Lasso error increases again. At the local minimum, Lasso provides the best features given its underlying assumptions. We suggest the user to run regular Lasso, to identify Lasso’s optimal performance point and then to define the desired number of features on the right-hand side of that point.

For our NN approach, we show that a cell’s training labels do not have to be unique. This is especially useful to take advantage of training data generated from DistMap’s probabilistic mapping output. We demonstrate how to properly split training and validation data when non-unique (Figure 3a) but correlated (Figure 3b) training labels are used in order to prevent data leakage. We hope that our approach will be helpful in the active research field of Weak Supervision (Zhou, 2018) and as probabilistic training labels become more commonplace. A lot of research in weak supervision for NNs has been around logistic regression and we hope that our techniques will help facilitate more applications in linear regression.

Not all decisions were consistent between Lasso-TopX and NN feature-selection approaches. For instance, the number of features (# of genes from RNASeq data used) and training labels (max MCC versus 95%) used during training differ between the approaches. Therefore, any differences in performance (Fig 3) and feature stability (Sup Figure S3, Sup Figure S5) also reflect various decisions made during the pre-processing stages. This was intentional as our goal was not to determine if NN or Lasso-TopX was better using the exact same data, but which independent approach would allow us to best address the sub-challenges.

One attribute of our models that is worth pointing out is that both the NN and Lasso approach could have also been used to predict X, Y, and Z spatial locations directly (not just for feature selection) because they were already optimized to predict a cell’s spatial position. However, we chose to use DistMap for this step to make the location prediction step consistent between the NN and Lasso approach and because the challenge organizers wanted 10 different location predictions per cell. Leveraging DistMap’s probabilistic mapping approach, using the MCC values made the later portion convenient because we were able to rank all spatial bins using the value of the coefficients.

Furthermore, we show that extra caution should be taken when deciding which data to be used in the location prediction process. We illustrated that in the case where DistMap/MCC approach was employed to predict cells’ 3-D locations, the provided genes’ expression binarized table should not be used as it leads to overestimation of performance. We observed this behavior for all feature selection methods (Lasso-TopX, Neural Nets, Random) and all score metrics, one of our own and three generated by the challenge’s organizers. We believe that this is happening due to data leakage because the binarized table supplied to the contestants was generated using the full *Single cell RNA sequencing* and *Reference Database* datasets. We show quantitively that even if the data were split into training and testing, using this table during location prediction transferred information from the test data to training (through calculating DistMap parameters), leading to an overestimation of performance.

Lastly, while identifying non-insitu genes was not a focus of the competition, we show that our methods were able to identify non-insitu genes that also contain spatial information. We show that the Lasso-TopX and NN approaches both reported similar genes. Surprisingly, when focusing on the most stable genes, slightly more than half (15 or 29) were non-insitu genes (Figure 4, Supplementary table S1). We believe that these *D. melanogaster* genes would be good candidates for exploring in future work involving spatial information.

## References

Ambroise,C. and McLachlan,G.J. (2002) Selection bias in gene extraction on the basis of microarray gene-expression data. Proc. Natl. Acad. Sci. U. S. A., 99, 6562–6566.

Bellman,R.E. (1961) Adaptive Control Processes A Guided Tour Princeton Legacy Library.

Fowlkes,C.C. et al. (2008) A Quantitative Spatiotemporal Atlas of Gene Expression in the Drosophila Blastoderm. Cell, 133, 364–374.

Friedman,J. et al. (2010) Regularization Paths for Generalized Linear Models Via Coordiante Descent. JournalofStatisticalSoftware, 33.

Gedeon,T.D. (1997) Data mining of inputs: analysing magnitude and functional measures. Int. J. Neural Syst., 8, 209–218.

Hahnioser,R.H.R. et al. (2000) Digital selection and analogue amplification coexist in a cortex-inspired silicon circuit. Nature, 405, 947–951.

Karaiskos,N. et al. (2017) The Drosophila embryo at single-cell transcriptome resolution. Science (80-.)., 358, 194–199.

Karathanasis,N. et al. (2014) Don’t use a cannon to kill the ? miRNA mosquito. Bioinformatics, 30, 1047–1048.

LeCun,Y. et al. (2015) Deep learning. Nature, 521, 436–444.

Luo,W. et al. (2016) Guidelines for developing and reporting machine learning predictive models in biomedical research: A multidisciplinary view. J. Med. Internet Res., 18, 1–10.

Nair,V. and Geoffrey E.,H. (2017) Rectified Linear Units Improve Restricted Boltzmann Machines. J. Appl. Biomech., 33, 384–387.

Paszke, Adam, Sam Gross, Soumith Chintala, Gregory Chanan, Edward Yang, Zachary DeVito, Zeming Lin, Alban Desmaison, Luca Antiga,A.L. (2017) Automatic differentiation in PyTorch. In, Conference on Neural Information Processing Systems (NIPS 2017). Long Beach, pp. 1–4.

Smialowski,P. et al. (2009) Pitfalls of supervised feature selection. Bioinformatics, 26, 440–443.

Srivastava,N. et al. (2014) Dropout: A Simple Way to Prevent Neural Networks from Overfitting. J. Mach. Learn. Res., 1929–1958.

Stolovitzky,G. et al. (2007) Dialogue on reverse-engineering assessment and methods: The DREAM of high-throughput pathway inference. Ann. N. Y. Acad. Sci., 1115, 1–22.

Tanevski,J. et al. (2019) Predicting cellular position in the Drosophila embryo from Single-Cell Transcriptomics data. bioRxiv.

Tibshirani,R. (1996) Regression Shrinkage and Selection via the Lasso. J. R. Stat. Soc. Ser. B, 267–288.

Verleysen,M. et al. (2003) On the effects of dimensionality on data analysis with neural networks. Lect. Notes Comput. Sci. (including Subser. Lect. Notes Artif. Intell. Lect. Notes Bioinformatics), 2687, 105–112.

Zeiler,M.D. (2012) ADADELTA: An Adaptive Learning Rate Method.

Zhou,Z.-H. (2018) A brief introduction to weakly supervised learning. Natl. Sci. Rev., 5, 44–53.

