## Supplementary Figures for "Machine Learning Approaches Identify Genes Containing Spatial Information from Single-Cell Transcriptomics Data"

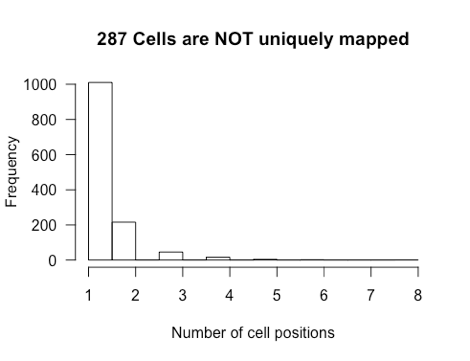

**Supplementary Figure S1***. DistMap (using all 84 inSitu genes) reports 287 cells that share their max-MCC position with another cell*

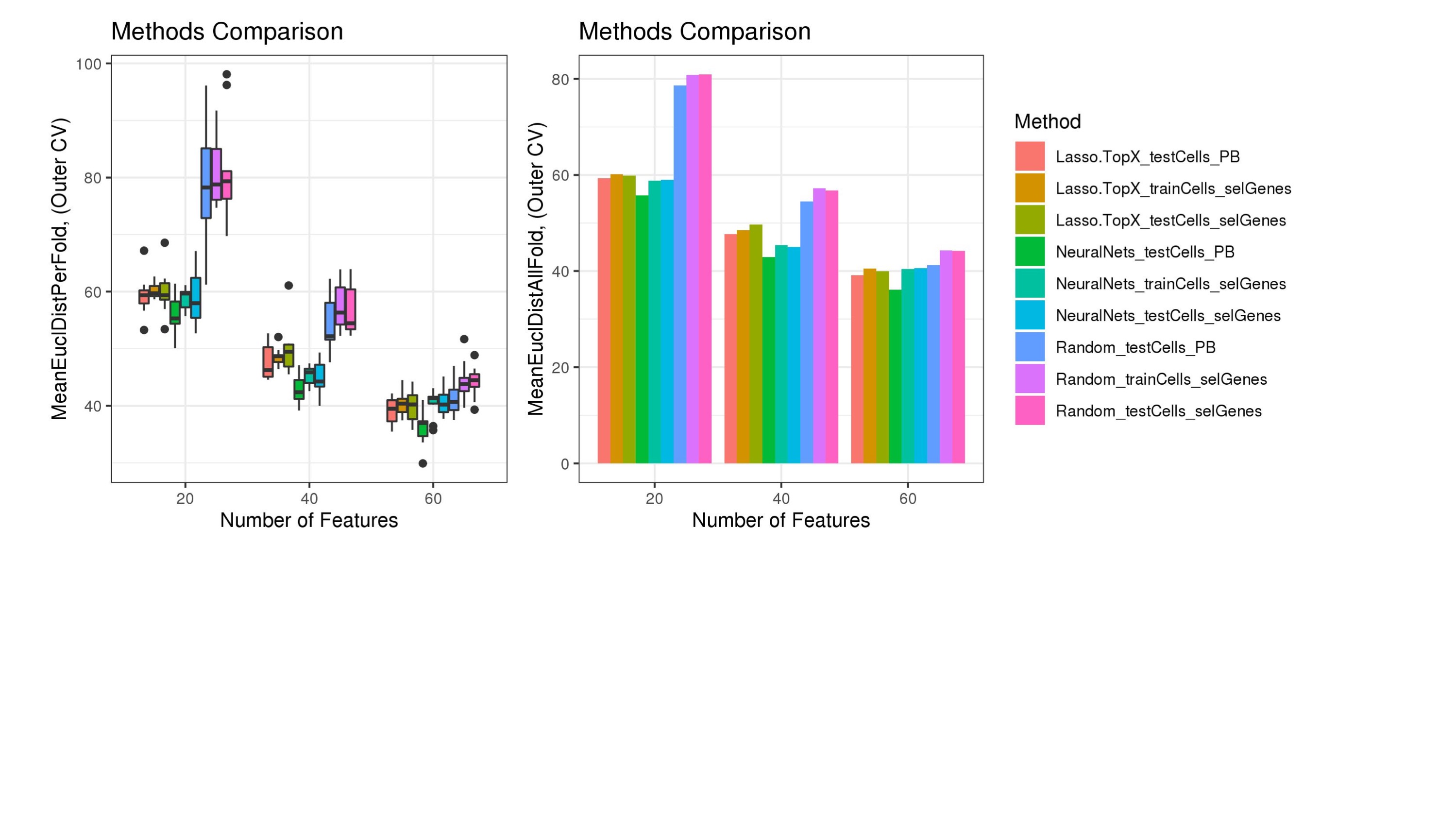

**Supplementary Figure S2***.* **Comparison of different methods – Blind metric.**

Lasso.TopX (Lasso.TopX_testCells_selGenes) performed better on selecting 60 features, sub-challenge 1, whereas NeuralNets (Light Blue - NeuralNets_testCells_selGenes) performed better in selecting 40 and 20 genes, sub-challenges 2 and 3. Training and testing errors present small differences which point to there is no overfitting taking place in either Lasso.TopX (Lasso.TopX_trainCells_selGenes and Lasso.TopX_testCells_selGenes) or NeuralNets (NeuralNets_trainsCells_selGenes and NeuralNets_testCells_selGenes). Both methods performed better than Random (Random_trainCells_selGenes and Random_testCells_selGenes). Using the provided binarized expression data, _PB extension, leaded to overestimation of performance accuracy, Lasso.TopX_testCells_PB, NeuralNets_testCells_PB, Random_testCells_PB. xaxis, different number of features (e.g. 60/40/20 for subchallenge #1/#2/#3 respectively). yaxis, left figure, MeanEuclDistPerFold, Mean Euclidean Distance Per outer cross validation fold. yaxis, right figure, MeanEuclDistAllFold, Mean Euclidean distance across all outer cross validation folds.

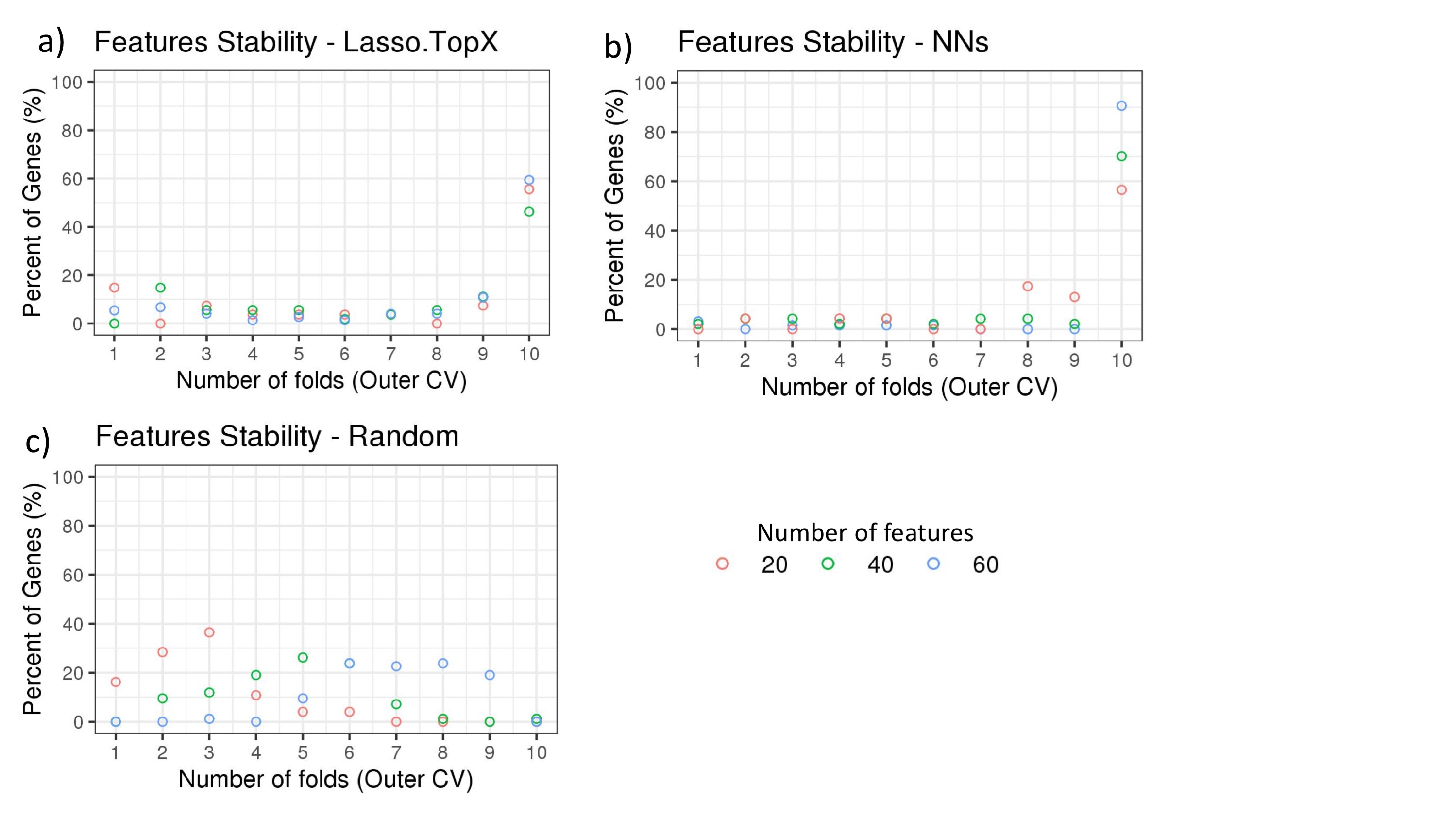

**Supplementary Figure S3. Features stability in Outer CV using inSitu genes.**

The percentage of common selected genes across the 10 fold outer cross validation is shown. a) In the Lasso.TopX case, 27 genes were selected across all folds for subchallenge 3, red dots. 4 genes or 15% percent were selected only in 1 fold, 2 genes or 7.4% percent were selected by 3 folds, etc. 15 or 56% percent were selected in all 10 folds. b) Similarly for the NNs case, out of a total of 47 genes selected across all folds for subchallenge 2, green dots, 33 genes or 70% percent were selected across all 10 folds of the outer cross validation. c) In the random case, 74 genes were selected across all folds for subchallenge 3, red dots. 12 genes or 16.2% percent were selected only in 1 fold, 21 genes or 28.4% were selected by two folds, etc. Importantly, for all subchallenges 0 genes were selected in all ten folds.

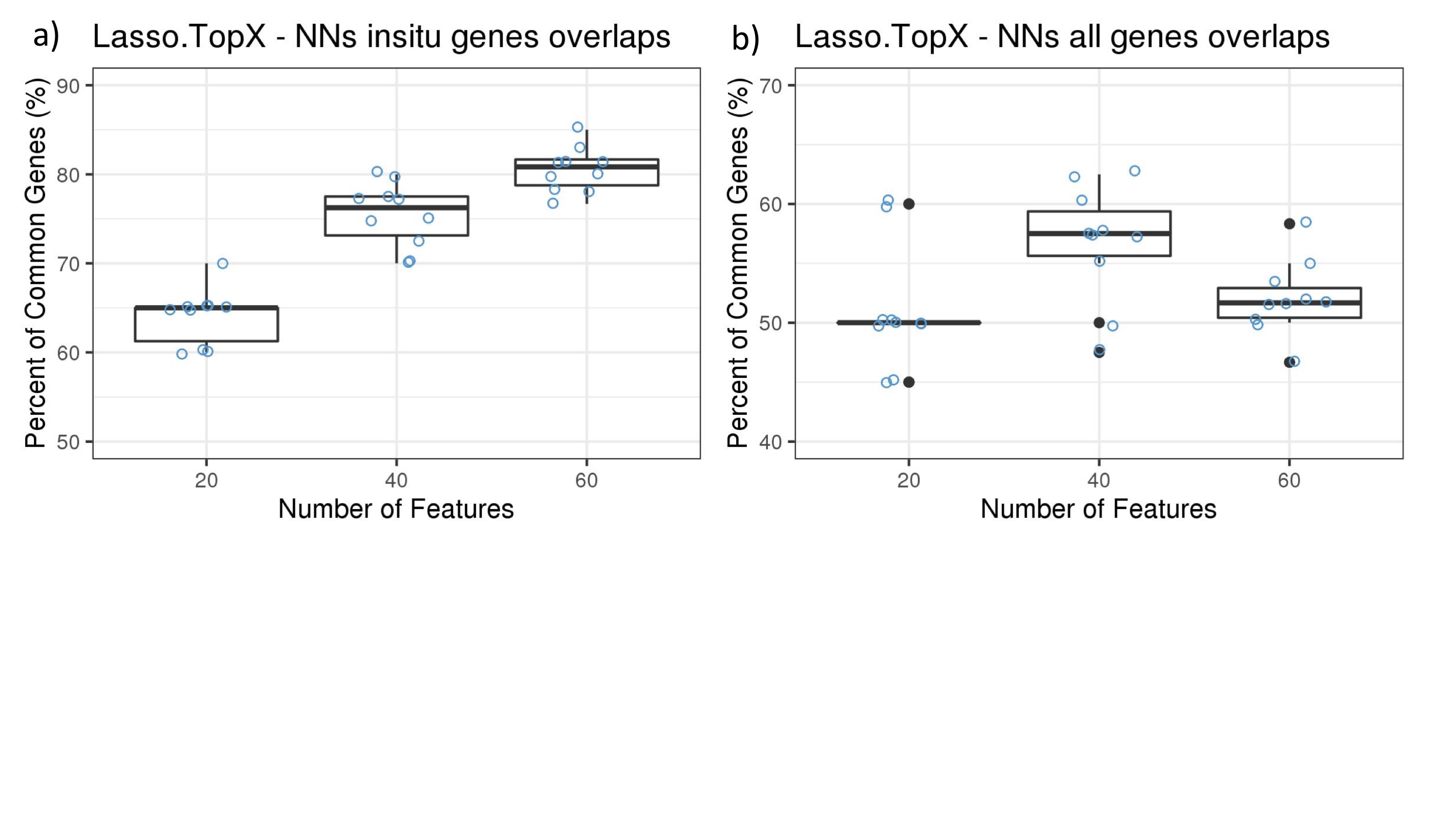

**Supplementary Figure S4.**

Lasso.TopX and NNs genes overlaps across the 10 fold outer cross validation, using either only inSitu genes,a, or all genes, b.

Yaxis: the percent of common genes between Neural Nets and Lasso.TopX, and

Xaxis: the number of features selected.

Boxplots and the corresponding data points in blue, are shown.

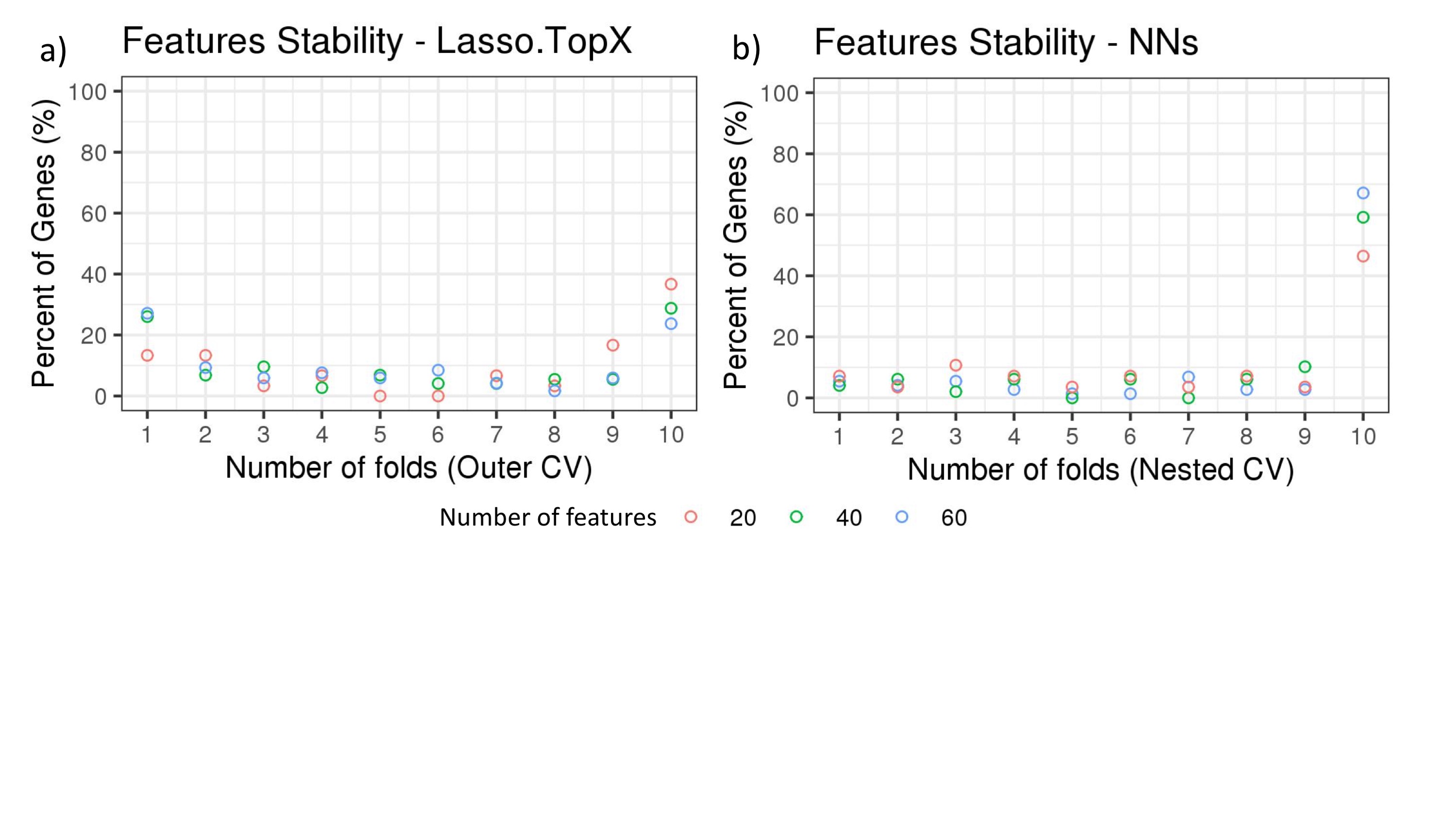

**Supplementary Figure S5. Features stability in Outer CV using all genes.**

The percentage of common selected genes across the 10 fold outer cross validation is shown. a) In the Lasso.TopX case, 30 genes were selected across all folds for subchallenge 3, red dots. 4 genes or 13.3% percent were selected only in 1 fold, 4 or 13.3% percent were selected by 3 folds, etc. 11 or 36.6% percent were selected in all 10 folds. b) In the NNs case, out of a total of 28 genes selected across all folds for subchallenge 3, red dots, 13 genes or 46.4% were selected across all 10 folds of the outer cross validation. Similar results are shown for subchallenge 1, and subchallenge 2, blue and green dots, respectively.
